# Gene expression profiling and protein-protein network analysis revealed prognostic hub biomarkers linking cancer risk in type 2 diabetic patients

**DOI:** 10.1101/2022.10.01.510254

**Authors:** Harshita Kasera, Rajveer Singh Shekhawat, Pankaj Yadav, Priyanka Singh

## Abstract

Type 2 diabetes mellitus (T2DM) and cancer are highly prevalent diseases imposing major health burden globally. Several epidemiological studies indicate increased susceptibility to cancer in T2DM patients. However, genetic factors linking T2DM with cancer are poorly studied so far. We used computational approach on the raw gene expression data of peripheral blood mononuclear cells of *Homo sapiens* available at the gene expression omnibus (GEO) database, to identify shared differentially expressed genes (DEGs) in T2DM and three common cancer types namely, pancreatic (PC), liver (LC) and breast cancer (BC). Additional functional and pathway enrichment analysis of identified common DEGs highlighted involvement of important biological pathways including cell cycle events, immune system process, cell morphogenesis, gene expression and metabolism. Furthermore, we retrieved the PPI network for crucial DEGs obtained from above analysis to deduce molecular level interactions. Based on the result of network analysis, we found 8, 5 and 9 common hub genes in T2DM *vs* PC, T2DM *vs* LC and T2DM *vs* BC, respectively. Overall, our analysis identified important genetic markers potentially able to predict the chances of pancreatic, liver and breast cancer onset in T2DM patients.

## Introduction

Type 2 diabetes mellitus (T2DM) is a highly prevalent metabolic disorder that can occur at any age albeit highly frequent in the middle (*i*.*e*., 45 years) to later age individuals. The onset of T2DM is marked by insulin resistance, a condition in which muscle, liver, and fat cells fail to use insulin properly. Eventually, the beta cells of pancreas cannot produce enough insulin either due to progressive beta cell mass reduction or their dysfunction. Thus, T2DM is characterized by insulin resistance and hyperglycemia in which blood glucose level increases[1]. The genome-wide association studies in the past have revealed some 403 distinct genetic variants in T2DM, which could influence beta-cell functioning, adipocytes, liver, skeletal muscle[2], and many other tissues. Therefore, it is not surprising that chronic T2DM condition could eventually result in further complications like nephropathy, cardiomyopathy, retinopathy, and neuropathy[3]. Consequentially, many differentially expressed genetic markers which could impart T2DM susceptibility were identified[4]. Subsequent bioinformatics analysis of these differentially expressed genes has revealed the genetic association of T2DM with several of the aforementioned complications[5]. These findings have advanced our understanding of complications arising due to T2DM and have prospective applications in designing personalized prognostic and diagnostic tools for such heterogenic human diseases.

Cancer is another heterogenic disease that is also the second most leading cause of human deaths[6]. Cancers are characterized usually by unrestricted growth of abnormal cells and, in some cases these abnormal cells could metastasize to other parts of the human body. Lung, prostate and breast cancers are among the most common cancer types[7]. It is well-known that T2DM and many common cancers share several risk factors like aging, obesity, and an unhealthy lifestyle[8]. Indeed, different epidemiological studies in the past suggest that T2DM condition increases the risk of several cancers, including liver, pancreatic[9], breast[10,11], and endometrial[12]. These epidemiological studies report standardized incidence ratios to indicate an increased risk of cancers in T2DM patients. Among the common cancer types, pancreatic and liver cancers showed the highest standardized incidence ratios in different populations of T2DM patients from Denmark, Tyrol/Austria, Taiwan, Sweden, Australia, the Chinese mainland[13], Finland[14], and Lithuania. In addition, a few meta-analysis studies reported an increased risk of breast cancer in diabetic women[10,11]. There is no clear molecular understanding of T2DM link to specific cancer types yet. However, the state of insulin resistance, hyperinsulinemia, hyperglycemia, chronic inflammation, and increased oxidative stress in T2DM could probably elicit mitogenic pathways and cause these cancers[15]. Despite the availability of extensive evidence from epidemiological and meta-analysis work linking cancer risk to T2DM, a systematic study of the shared genetic markers possibly predisposing this risk in T2DM patients is lacking for the common cancer types, namely pancreatic (PC), liver (LC), and breast (BC) cancer.

Herein, we perform gene expression analysis to identify predominant differential expressed genes (DEGs) in T2DM patients, posing a risk towards three common cancer types (PC, LC, and BC). We perform functional enrichment analysis of the identified shared DEGs to understand better an intricate interplay between T2DM and these cancer types. Moreover, we performed protein-protein interaction (PPI) network analysis to reveal the interactions among the shared genetic markers at the molecular level which allowed us to identify common hub genes shared by T2DM and three cancer types.

## Material & Methods

### Microarray Data Collection

The raw gene expression data of peripheral blood mononuclear cells (PBMCs) of *Homo sapiens* was available at the gene expression omnibus (GEO; URL: http://www.ncbi.nlm.nih.gov/geo/) database. All datasets used in this study were collected using the Affymetrix platform GPL570. **Table 1** provides a summary of different datasets used in the study. The <u>GSE15932</u> dataset comprised expression data of healthy individuals, T2DM, PC, and both T2DM and PC patients. The LC and BC gene expression data and their respective healthy controls were collected from the <u>GSE58208</u> and <u>GSE27562</u> datasets, respectively.

**Table 1.** Overview of GEO datasets used in this study. The dash (−) symbol indicates that the information is not available. Abbreviations: T2DM, type 2 diabetes mellitus; PC, pancreatic cancer; LC, liver cancer, BC, breast cancer

| Accession ID | Sample Type | Sample size | Gender | Age (in years) | Country |
| --- | --- | --- | --- | --- | --- |
| GSE15932 | T2DM | 8 | Male/Female | 43-80 | China |
|  | PC | 8 |  |  |  |
|  | T2DM & PC | 8 |  |  |  |
|  | Healthy | 8 |  |  |  |
| GSE58208 | LC | 10 | - | - | Singapore |
|  | Healthy | 5 |  |  |  |
|  | Others | 12 |  |  |  |
| GSE27562 | BC | 37 | Female | - | USA |
|  | Healthy | 31 |  |  |  |
|  | Others | 94 |  |  |  |

### Data Pre-processing and Identification of DEGs

The raw gene expression data was normalized using robust multichip average method available with affy package in the R software (version 4.0.1). The mean of log_2_ expression values were obtained for all genes in the respective datasets. Next, we calculated log_2_ fold change (log_2_FC) values for the all genes relative to their healthy controls. The prospective shared genetic markers between T2DM and three cancer types (PC, LC, and BC) were obtained by applying three filters. Our first filter is based on the p-value calculated using limma package in the R, indicating the statistical significance of differentially expressed genes. We considered genes having p-value 0.05 for further analysis. Our second filtering criteria is based on relative log_2_FC of gene expression, which is calculated for disease condition samples with respect to healthy condition samples indicating their biologically significant genes. We retained genes having absolute log_2_FC 0.5 for further analysis. Lastly, we retained correlated genes falling within the 10% interval from a regression line passing through origin (*i*.*e*., x=0 and y=0 on an xy plane) between log_2_FC in T2DM and respective cancer types. Only genes which were either upregulated or downregulated in both the conditions were selected for further analysis. Pearson correlation was calculated for the genes narrowed down after applying filtering criteria in each condition, using the cor function in the R software.

### Functional Enrichment Analysis of DEGs

The gene ontology (GO) enrichment analysis and Kyoto Encyclopedia of Genes and Genomes (KEGG) pathway analysis were performed to annotate the biological function of the DEGs using the online software GENECODIS. We considered a cut-off of false discovery rate (FDR) at 0.05 to define the significance level.

### Protein–Protein Interaction (PPI) Network Construction and Visualization

The PPI network analysis was performed based on the Search Tool for the Retrieval of Interacting Genes (STRING, URL: https://string-db.org), a database of known and predicted protein-protein interactions. To construct the PPI network, we used genes that were differentially expressed in both T2DM and three cancer types (PC, LC and BC). Interaction with a score > 0.8 was deemed statistically significant. The PPI network was constructed and visualized using the Cytoscape (version 3.8.2) which is an open-source software platform for visualizing molecular interaction networks and biological pathways.

### Hub Genes Identification

The hub genes were explored using cytoHubba application in the Cytoscape tool. For this purpose, the PPI network was analyzed to compute various topological features including degree, maximal clique, centrality, density of maximum neighborhood component, maximum neighborhood component, edge percolated component, bottleneck, eccentricity, closeness, radiality, betweenness, and stress. The top 20 nodes were considered as notable genes in the network for each of the computed topological features. The nodes which were common to all topological features were regarded as the important hub genes or key nodes in the network.

## Results

### Shared DEGs in T2DM and Three Common Cancer Types

We employed a three-tiered filtering criterion to identify the shared DEGs between T2DM and three cancer types namely, PC, LC and BC (**Figure 1**). The raw gene expression datasets were normalized for each GEO study using the robust multichip average method (**Figure S1**). We identified 113 common genes in T2DM *vs* PC comparison (**Figure 2a, S2a**). Of these, 81 were upregulated (**Figure S3a**) and 32 were downregulated (**Figure S3d**). Similarly, for T2DM *vs* LC comparison, we identified 275 common DEGs (**Figure 2b and S3b**), including 95 upregulated (**Figure S3b**) and 180 downregulated genes (**Figure S3e**). For T2DM *vs* BC comparison, we identified 26 common DEGs (**Figure 2c and S2c**), including 12 upregulated (**Figure S3c**) and 14 downregulated (**Figure S3f**). We observed strong Pearson’s correlation (0.98) for the identified shared DEGs between T2DM and three common cancer types.

**Figure 1.**
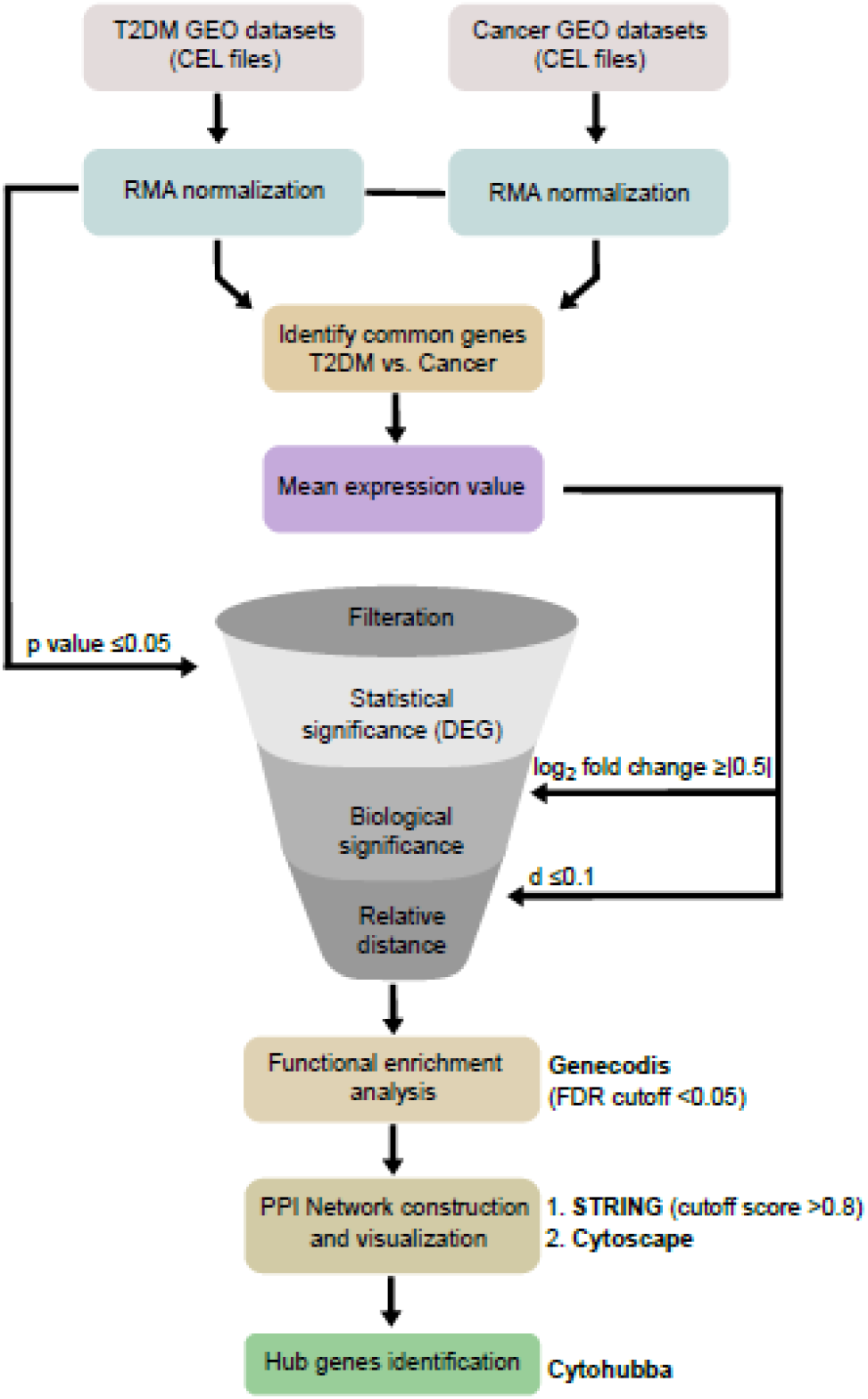
Schematic representation of the workflow used to identify differentially expressed genes (DEGs) and narrow down to the common hub genes in T2DM *vs* cancers datasets.

**Figure 2.**
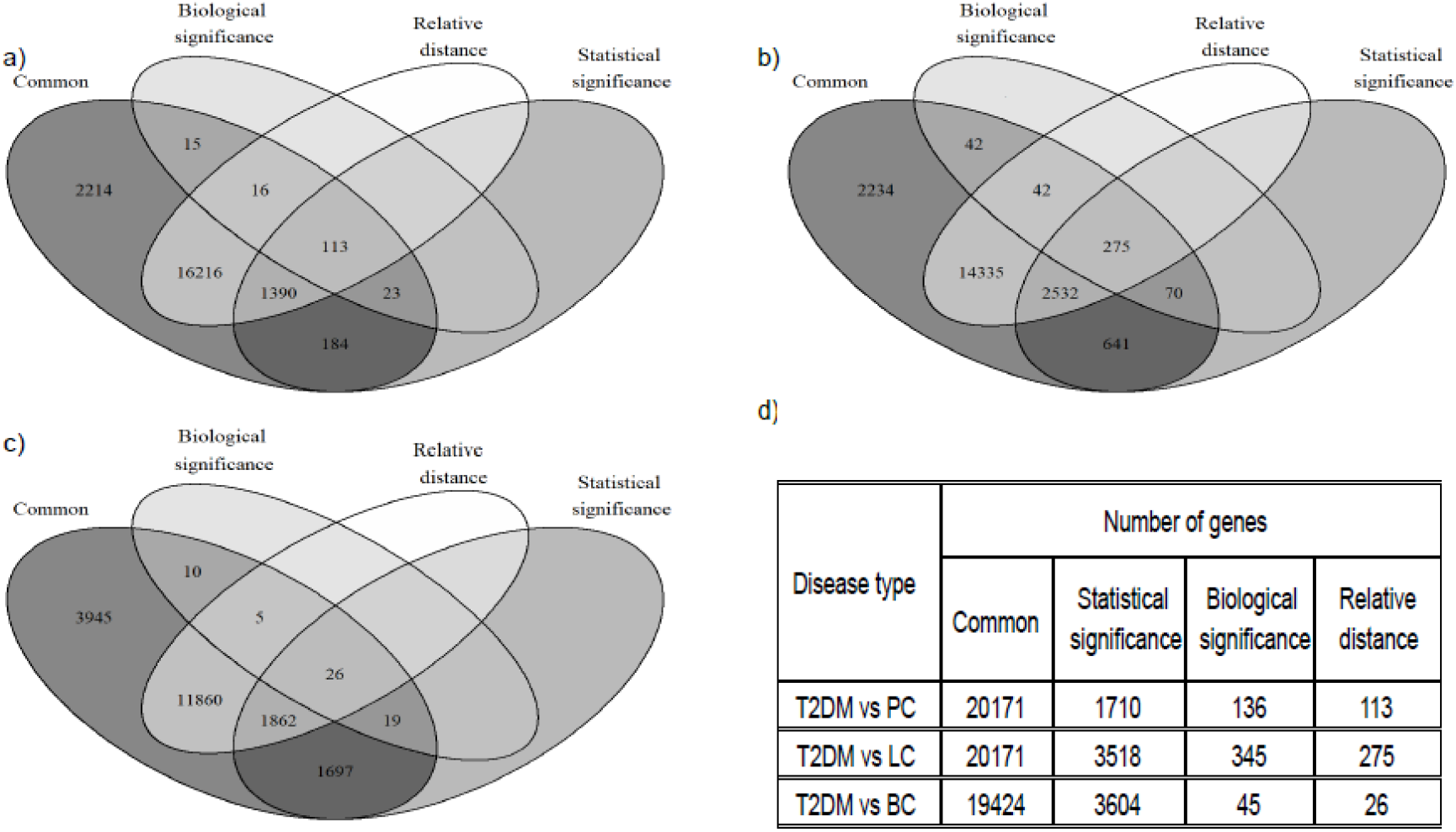
Venn diagram showing number of genes filtered with three-tiered filtering criteria (statistical significance (filter 1), biologically significance (filter 2) and 10% relative difference around linear regression line of correlated gene (filter 3). (a) 20,171 common genes obtained from GSE15932 dataset samples of T2DM and PC were narrowed down to 113 genes after the application of mentioned three filters. (b) Venn diagram representation of 20,171 common genes from GSE58208 dataset samples of T2DM and LC which can be narrowed down to 275 genes after applying three filtering criterion. (c) In case of T2DM *vs* BC 19,424 common gene genes from GSE27562 dataset samples could be narrowed down to 26 genes after applying filtering criterion. (d) Table summarizing the number of genes obtained after every filtering criterion in this study.

### Functional Enrichment and Pathway Analysis of Shared DEGs

In order to understand the functional relevance of identified common genes between T2DM and three cancers (PC, LC and BC), we utilized GO biological process and KEGG signaling pathway analysis tools. The KEGG signaling pathway analysis of common 113 DEGs in T2DM *vs* PC showed enrichment of metabolic pathways, neutrophil extracellular trap formation, viral carcinogenesis, cellular senescence, regulation of actin cytoskeleton, and transcriptional misregulation (**Table 2, section T2DM vs PC**). The GO analysis in **Table S1, section T2DM *vs* PC** showed significant enrichment of neutrophil degranulation (GO:0043312, p= 6.2×10^−13^), immune system process (GO:0002376, p=1.5×10^−3^), organelle localization by membrane tethering (GO:0140056, p=1.7×10^−3^), endoplasmic reticulum to vesicle mediated transport (GO:0006888, p=1.8×10^−3^), and cytokine mediated signaling pathway (GO:0019221, p=1.8×10^−3^).

**Table 2.** Shows top significantly enriched KEGG pathways involving the identified DEGs for three comparison T2DM *vs* PC, T2DM *vs* LC, and T2DM *vs* BC.

| Description | Annotation ID | Count | FDR | DEGs |
| --- | --- | --- | --- | --- |
| **T2DM vs PC** |  |  |  |  |
| Metabolic pathways | hsa01100 | 13 | $4.95 \times 10^{-9}$ | MGAT4B, TKT, RENBP, PDE7A, ND6, MGST1, ARG2, ACADVL, ALPL, FUT7, PHOSPHO1, NT5M, COMT |
| Neutrophil extracellular trap formation | hsa04613 | 5 | $3.91 \times 10^{-5}$ | H2BC21, H2AC18, VDAC3, RAC1, ITGB2 |
| Pathogenic *Escherichia coli* infection | hsa05130 | 5 | $7.69 \times 10^{-5}$ | RAC1, PAK2, OCLN, BRK1, PYCARD |
| Viral carcinogenesis | hsa05203 | 5 | $8.27 \times 10^{-5}$ | CCND3, H2BC21, VDAC3, REL, RAC1 |
| Cellular senescence | hsa04218 | 5 | $9.35 \times 10^{-5}$ | CCND3, CALM3, VDAC3, ZFP36L2, IGFBP3 |
| Pertussis | hsa05133 | 4 | $9.92 \times 10^{-5}$ | CD14, CALM3, ITGB2, PYCARD |
| Regulation of actin cytoskeleton | hsa04810 | 5 | $1.41 \times 10^{-4}$ | RAC1, PAK2, ITGB2, FGD3, BRK1 |
| Transcriptional misregulation in cancer | hsa05202 | 4 | $1.09 \times 10^{-3}$ | CEBPA, CD14, REL, IGFBP3 |
| Legionellosis | hsa05134 | 3 | $1.32 \times 10^{-3}$ | CD14, ITGB2, PYCARD |
| Ribosome | hsa03010 | 2 | $1.57 \times 10^{-3}$ | RPL38, RPL27A |
| **T2DM vs LC** |  |  |  |  |
| Metabolic pathways | hsa01100 | 30 | $5.81 \times 10^{-21}$ | SACM1L, TCIRG1, CDA, B4GALT5, ATP6V0D1, SUCLA2, ALDH5A1, UMPS, TKT, ATP6AP1, PKM, OXCT1, ARG2, IMPA2, ACADM, PHOSPHO2, HK3, GAA, ALOX5, ALDOA, GNPDA2, CBR4, AGL, ALG13, NADK, DLD, BDH2, AGPAT5, QRSL1, HIBCH |
| Herpes simplex virus 1 infection | hsa05168 | 12 | $8.03 \times 10^{-11}$ | ZNF195, ZNF184, ZNF84, SRSF7, CFP, BIRC3, ZNF879, ZNF320, ZNF420, ZNF573, ZNF566, PILRA |
| RNA transport | hsa03013 | 8 | $1.07 \times 10^{-9}$ | NUP160, NCBP2, RANBP2, NUP43, NUP35, EIF1AX, NUP133, TRNT1 |
| Phagosome | hsa04145 | 8 | $1.54 \times 10^{-7}$ | CORO1A, TCIRG1, ATP6V0D1, MARCO, TUBA4A, ATP6AP1, ITGB2, ITGAV |
| Human papillomavirus infection | hsa05165 | 10 | $4.10 \times 10^{-7}$ | TCIRG1, CCND3, ATP6V0D1, ATR, ATP6AP1, PKM, ITGAV, ITGA4, GSK3B, DLG1 |
| Fanconi anemia pathway | hsa03460 | 5 | $6.91 \times 10^{-7}$ | POLI, ATR, EME1, FANCF, POLK |
| Carbon metabolism | hsa01200 | 7 | $1.07 \times 10^{-6}$ | SUCLA2, TKT, PKM, HK3, ALDOA, DLD, HIBCH |
| Tuberculosis | hsa05152 | 8 | $2.12 \times 10^{-6}$ | CORO1A, TCIRG1, ATP6V0D1, CAMK2G, CAMK2D, ATP6AP1, NFYB, ITGB2 |
| Lysosome | hsa04142 | 6 | $2.17 \times 10^{-6}$ | TCIRG1, ATP6V0D1, AP1M1, ATP6AP1, GAA, MFSD8 |
| Ubiquitin mediated proteolysis | hsa04120 | 6 | $9.91 \times 10^{-6}$ | HERC4, WWP2, UBA2, CDC23, BIRC3, UBE2Q2 |
| **T2DM vs BC** |  |  |  |  |
| Ubiquitin mediated proteolysis | hsa04120 | 2 | $1.13 \times 10^{-2}$ | WWP2, UBE2M |
| Protein export | hsa03060 | 1 | $2.00 \times 10^{-2}$ | IMMP1L |
| Sulfur metabolism | hsa00920 | 1 | $2.00 \times 10^{-2}$ | SELENBP1 |
| Amyotrophic | hsa05014 | 2 | $2.17 \times 10^{-2}$ | ALYREF, ND6 |
| lateral sclerosis |  |  |  |  |
| mRNA surveillance pathway | hsa03015 | 1 | $4.56 \times 10^{-2}$ | ALYREF |
| Spliceosome | hsa03040 | 1 | $4.56 \times 10^{-2}$ | ALYREF |
| Osteoclast differentiation | hsa04380 | 1 | $4.98 \times 10^{-2}$ | JUNB |

In the T2DM *vs* LC comparison, the KEGG analysis of shared DEGs revealed enrichment in metabolic pathways, RNA transport, Fanconi anemia pathway, carbon metabolism (**Table 2, section T2DM *vs* LC**). We also found enrichment of GO biological processes shown in **Table S1, section T2DM *vs* LC**, which including regulation of transcription by RNA polymerase II (GO:0006357, p=2.7×10^−15^), regulation of cellular response to heat (GO:1900034, p=7.3×10^−13^), regulation of transcription (GO:0006355, p=7.6×10^−11^), neutrophil degranulation (GO:0043312, p=2.9×10^−10^), phosphorylation (GO:0016310, p=1.4×10^−8^), viral process (GO:0016032, p=5.1×10^−8^) and protein transport (GO:0015031, p=2.2×10^−6^).

Similarly, analysis of common DEGs in T2DM *vs* BC comparison led to the enrichment of ubiquitin mediated proteolysis, protein export, mRNA surveillance pathway, and spliceosome from KEGG (**Table 2, section T2DM *vs* BC**). The GO analysis shown in **Table S1, section T2DM *vs* BC** indicate significant enrichment of cell cycle arrest (GO:0007050, p=2.4×10^−3^), regulation of lymphocyte apoptotic process (GO:0070228, p=9.3×10^−3^), positive regulation of translational initiation in response to stress (GO:0032058, p=9.3×10^−3^) and cell morphogenesis (GO:0000902, p=9.7×10^−3^). The network cluster was obtained from GENECODIS software for an illustrative view of interconnection between functionally enriched pathways and identified DEGs as shown in **Figure S4**. The results of pathway enrichment analysis motivated us to further perform the interaction analysis at the molecular level.

### Deducing Molecular Level Interactions using PPI Network

The above analysis linked identified common genes to certain biological pathways which intrigued us to investigate their relationship at the molecular level. We constructed the PPI network maps of identified common gene between T2DM and three cancer types (PC, LC and BC). Our analysis yielded 35 nodes (genes) in T2DM *vs* PC, 112 nodes in T2DM *vs* LC, and 9 nodes in T2DM *vs* BC (**Table S2**). A close look at the PPI network indicated majority of upregulated nodes and a very few down-regulated nodes in the case of T2DM *vs* PC and T2DM *vs* BC (**Figure 3a and 3d**, respectively). For T2DM *vs* LC network, the number of down-regulated and up-regulated nodes were nearly the same (**Figure 3c)**.

In the PPI network, the node size reflects their degree and node color indicate the expression pattern, thus making it possible to deduce certain hub genes for respective conditions. Moreover, network analysis provides information on the enriched pathways involving major hub nodes as represented by a colored ring around each node (**Figure 3a and 3c**, respectively for T2DM *vs* PC and T2DM *vs* LC). This analysis revealed certain important genes which were associated with more than one biological pathway thus suggesting their evident involvement in the respective disease conditions. The important genes included, *CD14*, *CST3*, *ITGB2*, *PAK2*, *PYCARD* and *RAC1* from T2DM *vs* PC network and *NCBP2*, *NUP160*, *RANBP2*, *NUP43*, *NUP133*, *NUP35*, and *SRSF7* from T2DM *vs* LC network. We used these findings as a basis for further investigation of important hub genes.

**Figure 3.**
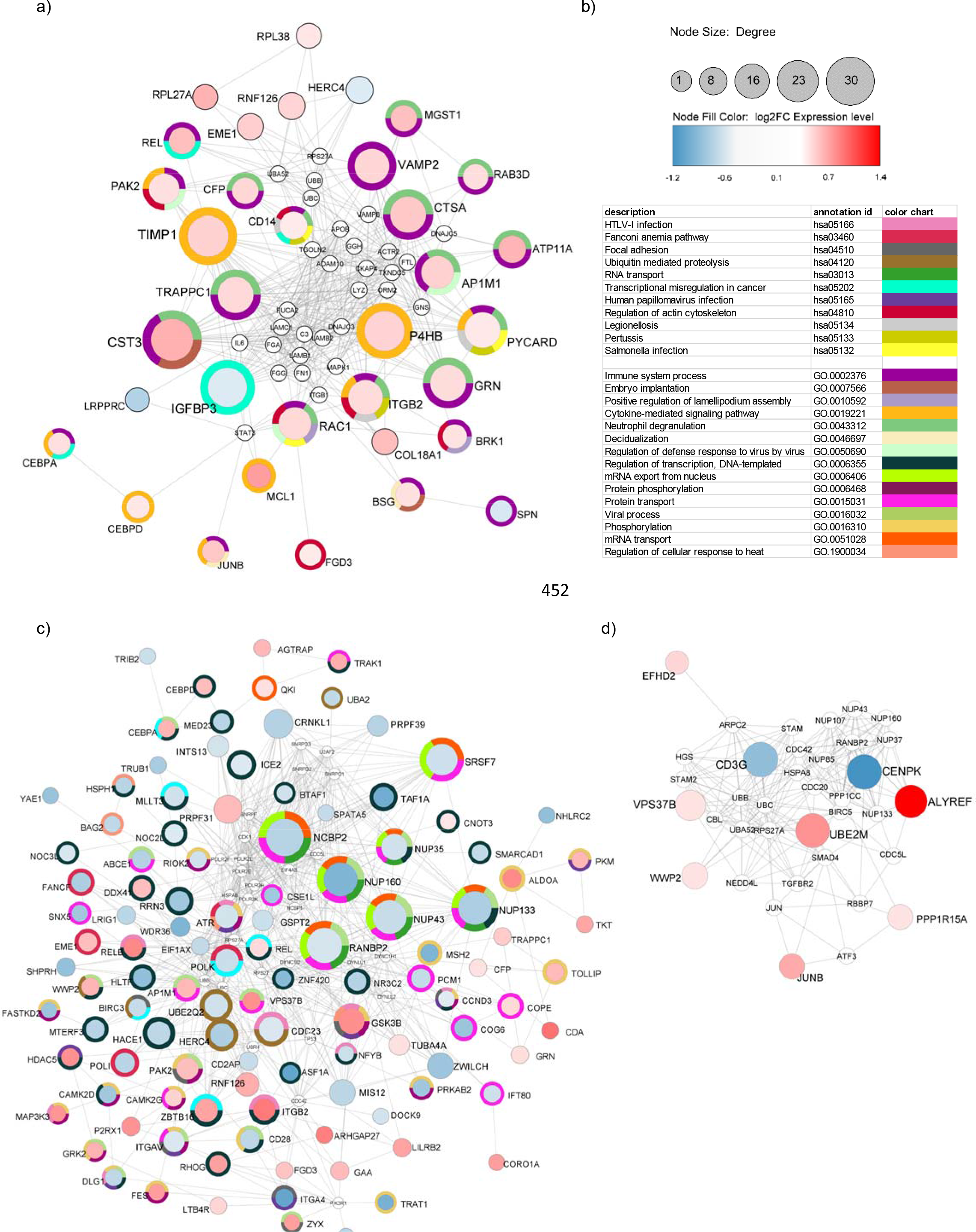
Protein-protein interaction network for (a) T2DM v*s* PC, (c) T2DM *vs* LC, (d) T2DM *vs* BC. The colored circle around the nodes represents different enriched pathways to which these nodes are linked. The small nodes (in white color) indicate additional interactors. The node color indicates overexpressed genes (in red) and under-expressed genes (in blue). In panel (b), node size represents the degree of the nodes and color intensity of the nodes represents the log_2_ fold change (FC) value of differentially expressed genes (DEGs). The color chart represents different enriched pathways along with their annotation identity.

### Hub Genes Identification and Analysis

We ranked the nodes in our PPI network analysis based up on eleven different topological features using cytoHubba plugin in the Cytoscape tool (see details in materials and methods section). This led us to identification of the top 20 genes for both T2DM *vs* PC and T2DM *vs* LC (**Table S3**). We noticed several hub genes including *CST3*, *TIMP1*, *P4HB* and *NCBP2*, *RANBP2*, *NUP133* that were top-ranked in almost all the computed topological features for both T2DM *vs* PC and T2DM *vs* LC. This analysis was not performed for T2DM *vs* BC due to insufficient number of identified common genes. We further narrowed down common hub genes based on their top ranking and commonness across the computed topological features. Accordingly, we could identify 8 genes as the hub genes for T2DM *vs* PC comparison. The relative log_2_ fold expression of these genes in two diseased conditions is represented in **figure 4a and 4c**. For T2DM *vs* LC comparison, we identified 5 hub genes. We found close expression of these genes in T2DM and LC disease condition as indicated in the **figure 4d**. Similarly, for T2DM *vs* BC comparison, 9 hub genes were identified as represented in the **figure 4b and 4e**. Overall, we identified 22 hub genes considering the three comparisons which could serve as potential biomarkers.

**Figure 4.**
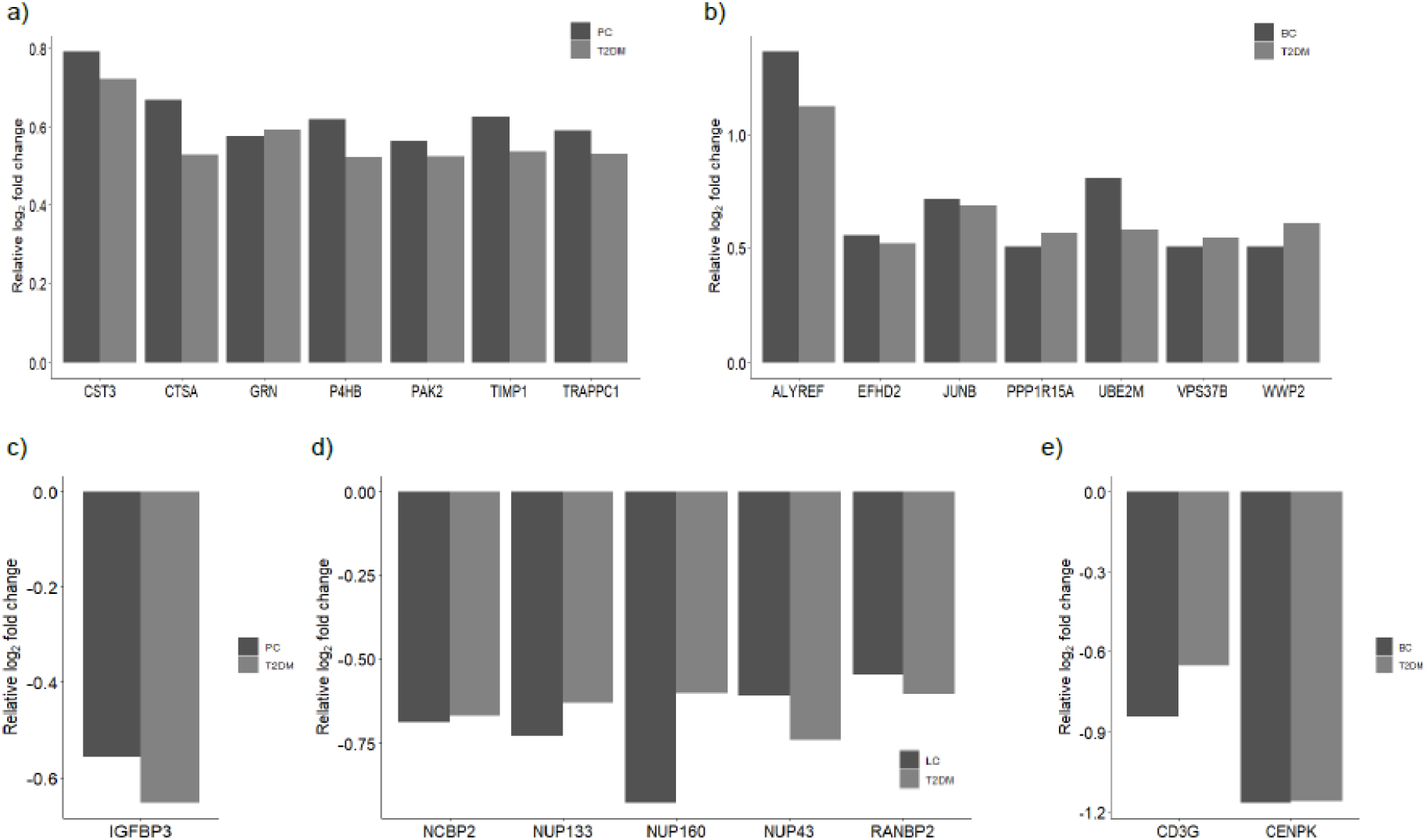
The barplot shows log_2_ fold change expression of identified hub genes in both disease conditions. The upregulated genes are shown for (a) T2DM *vs* PC and (b) T2DM *vs* BC. The downregulated genes are shown for (c) T2DM *vs* PC, (d) T2DM *vs* LC and (e) T2DM *vs* BC.

## Discussion

T2DM and cancer are widespread globally and have burdened the health sector throughout the world. In recent past, several epidemiological studies have indicated a causal link between T2DM and common cancer types. However, the genetic association of T2DM to these cancers remains largely unknown. Our work identifies shared genetic association between T2DM and three common cancer types, namely PC, LC and BC. Our analysis identified 113, 275, and 26 genes showing an overlap between T2DM and three cancers *i*.*e*., PC, LC and BC, respectively. Interestingly, we found a 71% (80 out of 113 genes) overlap between our identified list of correlated genes in the case of T2DM vs PC by comparing them with an independent GEO dataset comprising patients with both conditions (**Figure S5**). We could not perform similar validation for the T2DM *vs* LC and T2DM *vs* BC due to unavailability of such relevant studies.

Our analysis of KEGG pathway and GO biological process associated with identified common gene in T2DM *vs* PC and T2DM *vs* LC showed enrichment of metabolic pathway or carbon metabolism[16] which is expected specially because T2DM is a metabolic disorder and pancreas and liver are closely associated with metabolism. The neutrophil extracellular trap formation found to be enriched in T2DM *vs* PC and degranulation enriched in T2DM *vs* PC and T2DM *vs* LC are known to be associated with diabetes[17] as well as cancer cell progression, metastasis and activating dormant cancer cell[18]. Several viral infections like Hepatitis B and Hepatitis C are known to cause liver cancer. These viral infections have also been investigated for possible linkage with the pancreatic cancer especially due to chronic local inflammation[19]. These could also possibly play a role in development of T2DM. Our analysis also identified common genes belonging to the immune system process, as any imbalance in immune system is directly correlated with tumor progression and hyperglycemia[20]. Furthermore, remodeling of actin cytoskeleton is observed during insulin secretion from the pancreatic β-cell and could also affect cancer cell migration[21]. Membrane trafficking process are required for glucose uptake induced by insulin and several membrane associated proteins also contribute to tumor progression[22]. Our analysis also revealed important pathways contribution to gene expression which include transcription, RNA transport, RNA polymerase II transcription, translation initiation, mRNA surveillance, phosphorylation, protein export, spliceosomes, Fanconi anemia pathway involved in DNA repair and ubiquitin-mediated proteolysis. Such regulatory networks of related genes are necessary for maintaining homeostasis, any alteration can lead to certain diseases and disorders[23]. Moreover, cytokine-mediated response which elicit immune cell response was found to be enriched in both T2DM *vs* PC and T2DM *vs* BC. The lymphocyte apoptosis is crucial to both diabetes and cancer as several studies suggest that diabetes can lead to lymphocyte apoptosis and in case of cancer this process is carried out as a protective mechanism to weaken the immune system and immune escaping[24]. Cellular senescence resulting in aging, a contributing factor in metabolic dysfunction, and inflammation which can lead to T2DM and cancer phenotype[25]. Cell morphogenesis is associated with epithelial plasticity and epithelial to mesenchymal transition (EMT) which are also vital in metastasis and progression of any cancer[26].

In nutshell, we identified 8 common hub genes (*CST3*, *CTSA*, *GRN*, *IGFBP3*, *P4HB*, *PAK2*, *TIMP1*, *TRAPPC1*) in T2DM *vs* PC, 5 common hub genes (*NCBP2*, *NUP133*, *NUP160*, *NUP43*, *RANBP2*) in T2DM *vs* LC and 9 common hub genes (*ALYREF*, *CD3G*, *CENPK*, *EFHD2*, *JUNB*, *PPP1R15A*, *UBE2M*, *VPS37B*, *WWP2*) in T2DM *vs* BC, respectively. The detailed function and description related to these 22 hub genes is provided in **Table S4**. CST3 (cystatin C) has been explored as a potential linkage marker in T2DM with associated complication[27]. CTSA (serine protease cathepsin A) and its members are involved in cell proliferation, migration, and apoptosis. Its family members cathepsin D and L have been shown to be impaired in T2DM patients[28]. GRN (granulin precursor) is known for its role in tumor proliferation, insulin resistance and adipokine related obesity[29]. IGFBP3 (insulin like growth factor binding protein 3) have potent insulin-antagonizing capability[30] which could cause T2DM and its disruption could also lead to pathophysiology in many cancers[31]. P4HB (Prolyl 4-hydroxylase, beta polypeptide) is an endoplasmic reticulum chaperone, required for proinsulin maturation[32] proposed to be involved in EMT and β-catenin/Snail pathway in cancer cells[33]. PAK2 (P21 (RAC1) Activated Kinase 2) is a serine/threonine protein kinase associated with signaling pathways having role in cell proliferation, homeostasis, inducing apoptosis, and impede cell cycle progression. TIMP1 (tissue inhibitor of matrix metalloproteinases-1) regulating extracellular matrix, angiogenesis and cell proliferation, is also associated with chronic inflammation of adipose tissue in T2DM patients. TRAPPC1 (Trafficking Protein Particle Complex Subunit 1) direct link to T2DM and cancer is not quite clear but its family members are known to be involved in MAP kinase ERK2 mediated downstream signaling[34]. Similarly, *VPS37B* (Vacuolar Protein Sorting-Associated Protein 37B) common hub gene in T2DM *vs* BC, code for protein involved in vesicle-mediate transport. Disruption of membrane trafficking pathways could lead to several human diseases[35]. NCBP2 is a nuclear cap-binding protein which can regulate the expression of small noncoding RNA in T2DM[36] and hepatocellular carcinoma[37]. NUP133, NUP160, NUP43, RANBP2 and ALYREF are nuclear pore associated proteins and they could affect gene expression both in T2DM and cancer[38]. *CD3G* code for CD3-T-cell surface glycoprotein which upon mis-regulation could elicit immune response and result into inflammation[39]. EFHD2 belongs to the family of EF-hand calcium binding proteins which is associated with metastatic activity of several cancer types[40]. We identified a number of common hub genes for T2DM *vs* LC and T2DM *vs* BC comparisions which are involved in cellular homeostasis, cell signaling and proliferation.

This includes CENPK, centromere associated protein; JUNB, a transcription factor; PPP1R15A (Protein Phosphatase 1 Regulatory Subunit 15A) regulating protein phosphorylation and protein turnover enzymes including UBE2M (Ubiquitin Conjugating Enzyme E2 M), and WWP2 (WW Domain Containing E3 Ubiquitin Protein Ligase 2).

Our analysis provides an insight towards different biological and molecular events which could possibly link T2DM with three common cancer types (PC, LC and BC). These identified genetic markers holds potential to predict the chances of cancer onset in T2DM patients. Importantly, such markers in T2DM patients predisposing increased cancer risk could help better diagnosis of cancers at early stages and also benefit developing personalized therapeutic strategies.

## Author Contributions

HK collected study data, performed primary DEG analysis and designed the working strategy; RSS performed functional enrichment, PPI network and hub gene analysis; PS conceptualized the work and together with PY designed and supervised the study; all authors contributed to writing the manuscript.

## Abbreviations

T2DM: Type 2 Diabetes Mellitus
PC: Pancreatic Cancer
LC: Lung Cancer
BC: Breast Cancer
DEGs: Differentially Expressed Genes
PPI: Protein-protein interaction
PBMC: Peripheral blood mononuclear cells
log_2_FC: log_2_ fold change
GO: Gene ontology
KEGG: Kyoto encyclopedia of genes and genomes
FDR: False discovery rate

## Acknowledgements and Funding

HK (file number:09/1125(0017)/2020-EMR-I) and RSS (file number:09/1125(0019)/2021-EMR-I) are supported by the CSIR-NET fellowship. PY acknowledges the support from seed grant (project number I/SEED/PY/20200037) funded by Indian Institute of Technology, Jodhpur, India. PS is thankful to Science and Engineering Research Board (ECR/2017/001410), Department of Biotechnology (BT/12/IYBA/2019/02) and Board of Research in Nuclear Sciences (55/14/02/2021-BRNS/10206) for the financial support.

## Conflict of Interest

Authors declare that there are no competing financial interests.

## Additional Information

The online version of this article contains a data supplement.

## Notes

### Competing Interest Statement

The authors have declared no competing interest.

