## Supplementary Data for "Gene expression profiling and protein-protein network analysis revealed prognostic hub biomarkers linking cancer risk in type 2 diabetic patients"

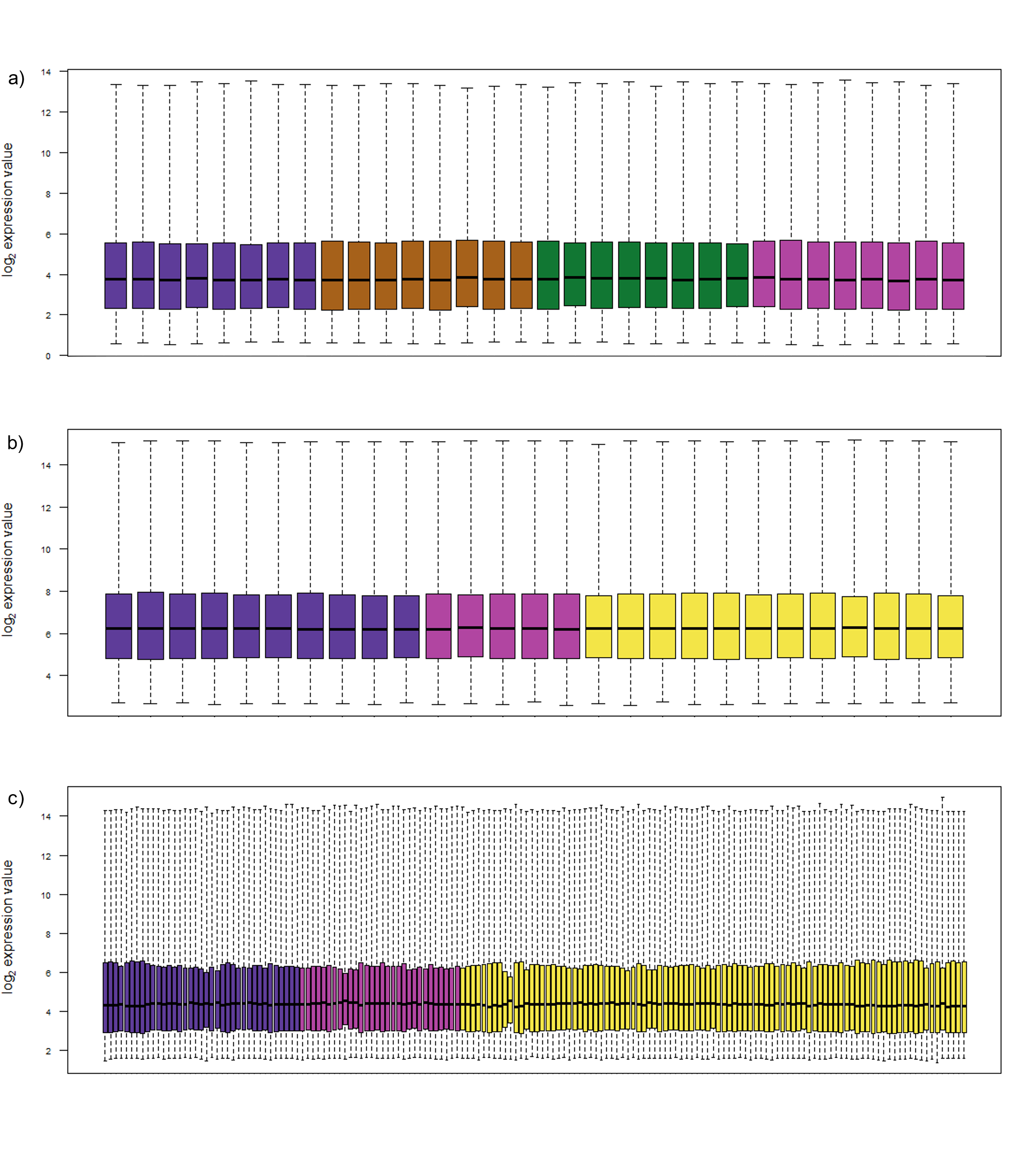

**Figure S1**. Boxplot of RMA (robust multichip average) normalized gene expression datasets: **(a)** GSE15932 study having patients suffering from only type 2 diabetes mellitus (T2DM) (in indigo), only pancreatic cancer (PC) (in brown), both T2DM and PC (in green) and healthy patients (in purple), **(b)** GSE58208 study having patients suffering from liver cancer (LC) (indigo), healthy patients (in purple) and others (in yellow). **(c)** GSE27562 study having patients suffering from breast cancer (BC) (in indigo), healthy patients (in purple) and others (in yellow).

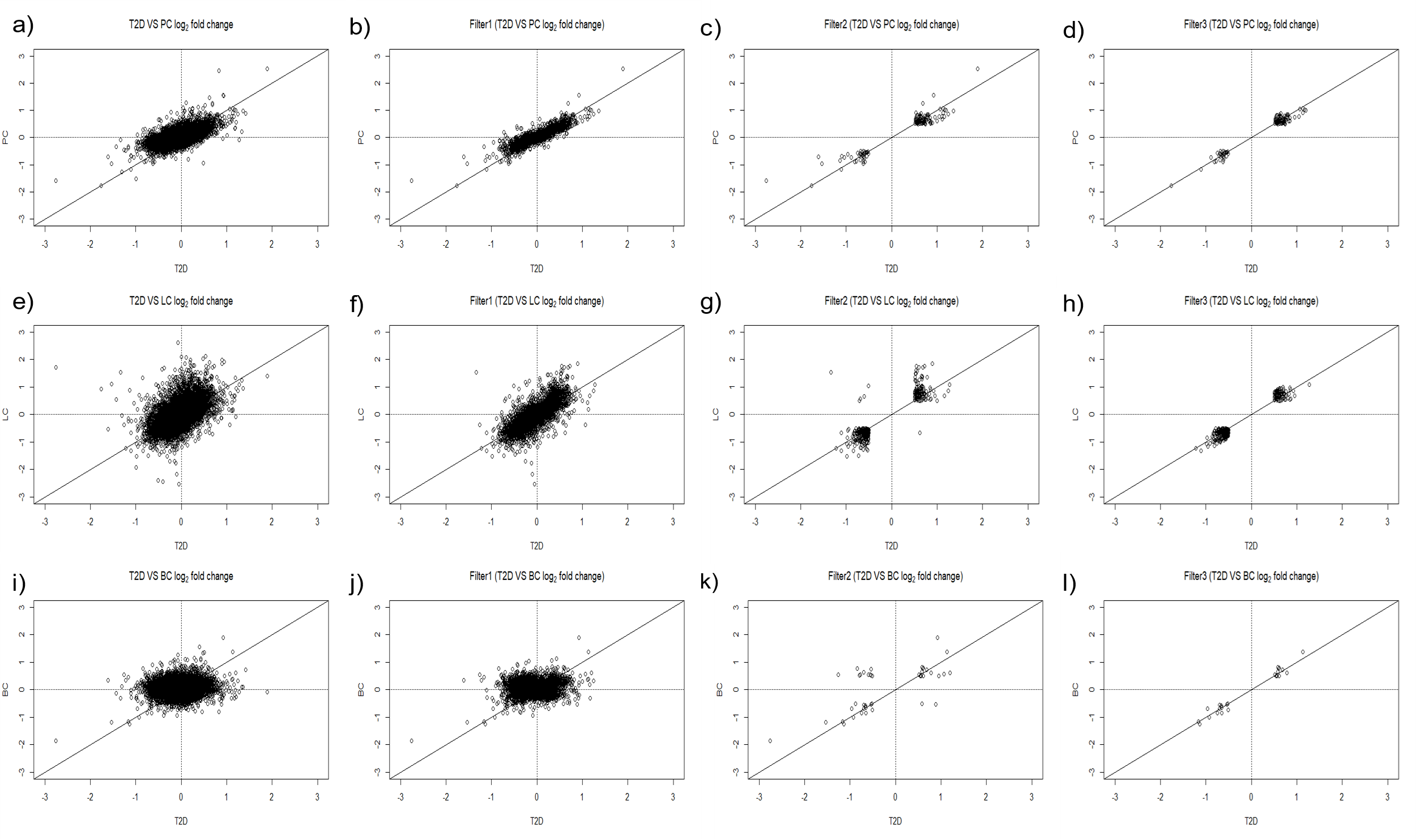

**Figure** **S2**. Scatter plot showing filtration of genes between T2DM and three cancer types (PC, LC, BC). **(a-d)** T2DM *vs* PC. Initially, expression data on 20,171 common genes was available (panel a). After applying filtering criterion, 113 genes were remained for further analysis. **(e-h)** T2DM *vs* LC. Initially, expression data on 20,171 common genes was available (panel e). 275 genes were obtained after the filtering criterion. **(i-l)** T2DM *vs* breast cancer (BC). 19424 genes expression data was provided in common and 26 genes were obtained after the filtering criterion

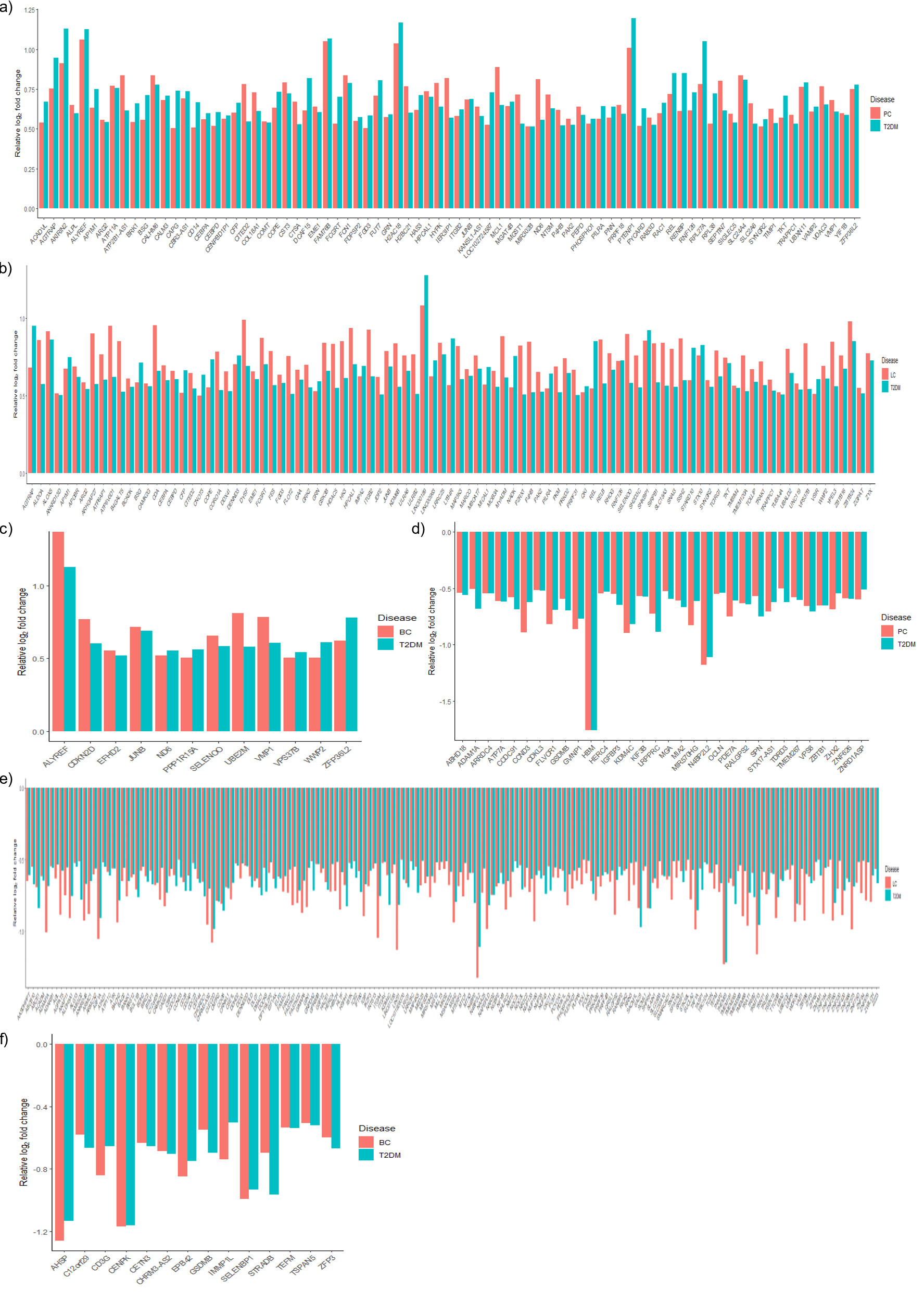

**Figure S3.** Expression pattern of the genes obtained after filtering in T2DM and cancer types. Upregulated genes having log_2_ fold change values greater than 0.5 **(a)** T2DM *vs* PC **(b)** T2DM *vs* LC **(c)** T2DM *vs* BC. Downregulated genes having log_2_ fold change lesser than -0.5 **(d)** T2DM *vs* PC, **(e)** T2DM *vs* LC, **(f)** T2DM *vs* BC

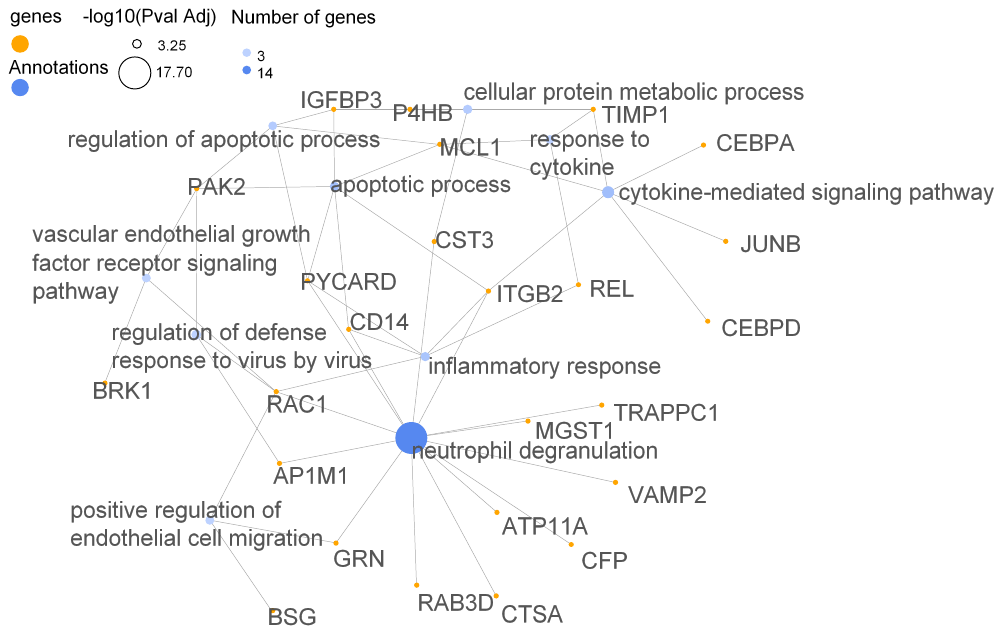

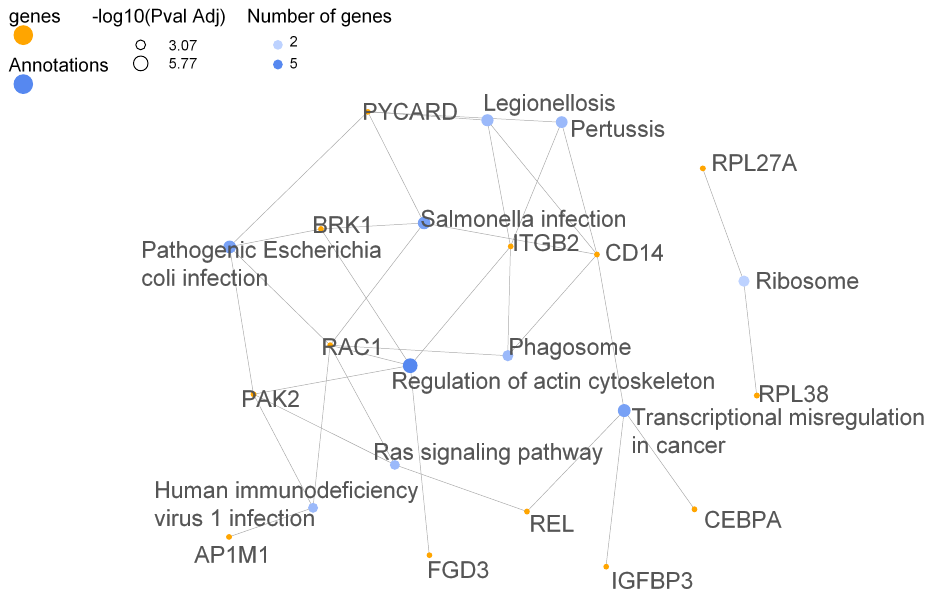

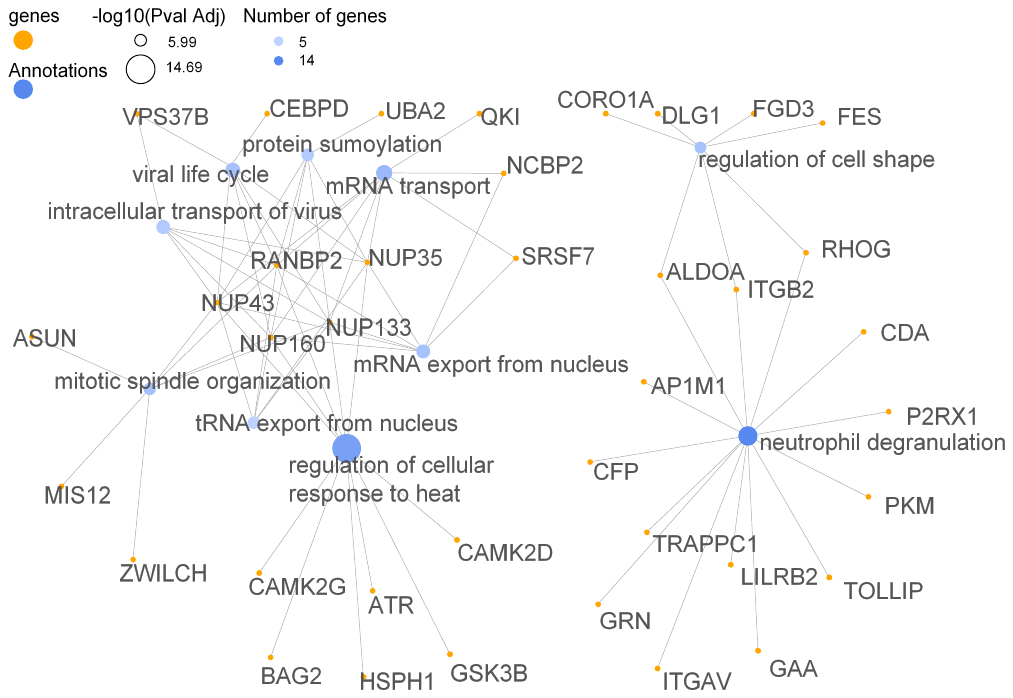

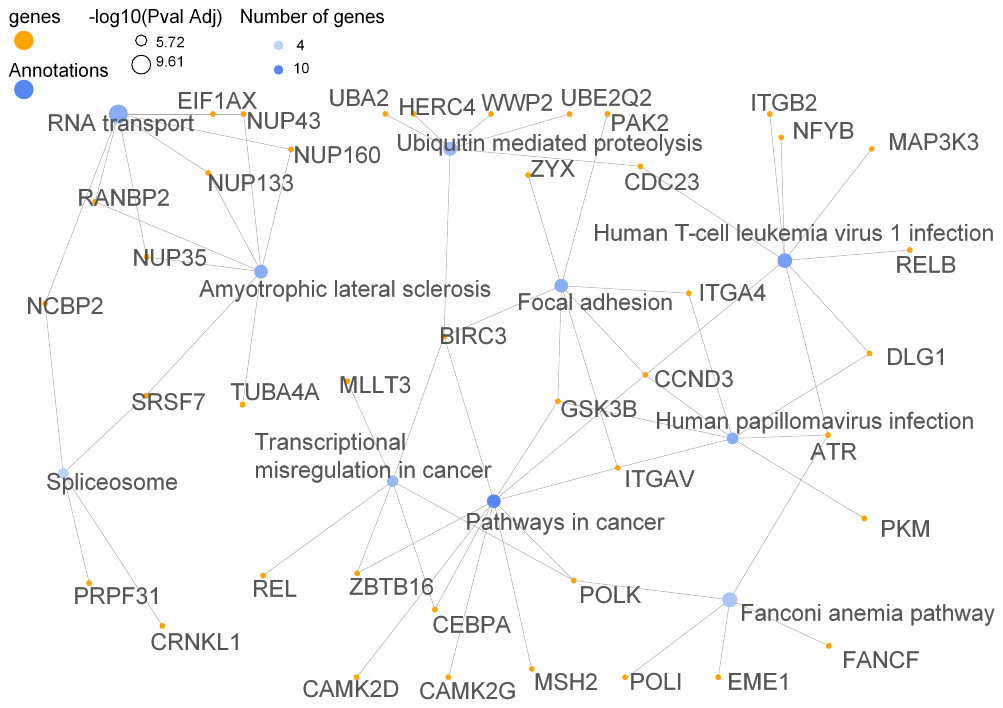

a)

b)

c)

d)

**Figure S4:** Shows the network of enriched pathways along with the DEGs associated with the pathways. KEGG pathway network for T2DM *vs* PC and T2DM *vs* LC (panel a and c, respectively). GO BP network for T2DM *vs* PC and T2DM *vs* LC (panel b and d, respectively).

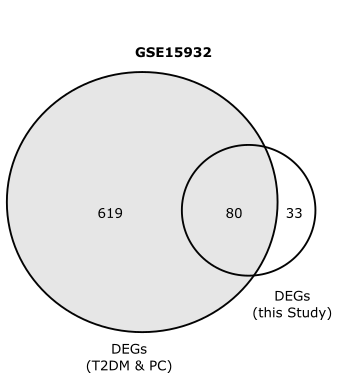

**Figure S5.** Venn diagram showing number of genes obtained from filtering in this study that are overlapping with differentially expressed genes (DEGs) in patients suffering from both T2DM and PC. Note that 71% (80 genes out of 113) of genes obtained from this study overlaps with DEG in patients suffering from T2DM and PC.

**Table S1**. Top significantly enriched gene ontology (GO) pathways for the shared differentially expressed genes from three comparison, namely T2DM *vs* PC, T2DM *vs* LC and T2DM *vs* BC.

| **Description** | **Annotation ID** | **Count** | **FDR** | | **Genes** |
| --- | --- | --- | --- | --- | --- |
| **T2DM *vs* PC** |  | | |  | |
| Neutrophil degranulation | GO:0043312 | 16 | 6.20x10^-13^ | | ATP11A,CD14,RAB3D,AP1M1,SIGLEC5,VAMP2,RAC1,CTSA,CFP,MGST1,ITGB2,GRN,FCN1,TRAPPC1,CST3,PYCARD |
| Positive regulation of lamellipodium assembly | GO:0010592 | 4 | 1.14x10^-4^ | | RAC1,ATP7A,OCLN,BRK1 |
| In utero embryonic development | GO:0001701 | 6 | 6.72x10^-4^ | | ATP11A,CITED2,ATP7A,JUNB,BRK1,FLVCR1 |
| Innate immune response | GO:0045087 | 8 | 1.09x10^-3^ | | ZBTB1,CD14,REL,CFP,ARG2,FCN1,AKIRIN2,PYCARD |
| Immune system process | GO:0002376 | 8 | 1.46x10^-3^ | | ZBTB1,CD14,CFP,ARG2,FCN1,DCAF15,AKIRIN2,PYCARD |
| Decidualization | GO:0046697 | 3 | 1.60x10^-3^ | | CITED2,BSG,JUNB |
| Organelle localization by membrane tethering | GO:0140056 | 2 | 1.69x10^-3^ | | CALM3,VMP1 |
| Viral process | GO:0016032 | 8 | 1.71x10^-3^ | | CEBPA,ALYREF,SYNGR2,AP1M1,CALM3,BSG,PAK2,PILRA |
| Endoplasmic reticulum to Golgi vesicle-mediated transport | GO:0006888 | 5 | 1.78x10^-3^ | | COPE,MIA2,YIF1B,TRAPPC1,IER3IP1 |
| Cytokine-mediated signaling pathway | GO:0019221 | 6 | 1.81x10^-3^ | | CEBPD,CEBPA,TIMP1,MCL1,JUNB,ITGB2 |
| **T2DM *vs* LC** |  |  |  | | |
| Regulation of transcription by RNA polymerase II | GO:0006357 | 34 | 2.69x10^-15^ | | KDM4C,TRAK1,IKZF3,CEBPD,CEBPA,ZBTB6,CITED2,ZBTB24,MED23,CAMK2D,ZNF195,ZNF184,ZBTB16,ZNF84,ZNF75D,RELB,REL,NFYB,NR3C2,JUNB,ZFP62,ZNF879,ZNF260,SNAI3,ZNF320,ZNF420,ZNF573,ZFP3,JDP2,ZNF566,ZGPAT,BCL11B,BACH2,ELP2 |
| Regulation of cellular response to heat | GO:1900034 | 12 | 7.34x10^-13^ | | NUP160,HSPH1,BAG2,CAMK2G,CAMK2D,RANBP2,ATR,GSK3B,NUP43,NUP35,NUP133,CHORDC1 |
| Regulation of transcription, DNA-templated | GO:0006355 | 27 | 7.62x10^-11^ | | CEBPD,CEBPA,CITED2,MED23,ZNF195,ZNF184,ZBTB16,ZNF84,ZNF75D,RELB,REL,CNOT3,NFYB,NR3C2,MLLT3,JUNB,ZNF879,SNAI3,ZNF320,ZNF420,NUP35,ZNF573,JDP2,ZNF566,BACH2,DDX1,MTERF3 |
| Neutrophil degranulation | GO:0043312 | 19 | 2.88x10^-10^ | | SIRPB1,TCIRG1,CDA,LILRB2,AP1M1,PKM,CFP,P2RX1,RHOG,ITGB2,ITGAV,HK3,GRN,GAA,ALOX5,ALDOA,AGL,TRAPPC1,TOLLIP |
| Phosphorylation | GO:0016310 | 20 | 1.41x10^-8^ | | PRKD2,STK38L,MAP4K5,BCKDK,PRPF4B,CAMK2G,CAMK2D,ATR,PKM,PAK2,MAP3K3,HK3,GSK3B,PDIK1L,FES,NADK,GRK2,RIOK2,N4BP2,TOLLIP |
| Viral process | GO:0016032 | 18 | 5.13x10^-8^ | | NUP160,WWP2,CEBPA,SYNGR2,AP1M1,ZYX,BSG,ABCE1,RANBP2,PAK2,ITGAV,NUP43,NUP35,EIF1AX,DLG1,DDX1,NUP133,PILRA |
| Intracellular transport of virus | GO:0075733 | 7 | 3.72x10^-7^ | | NUP160,NMT2,RANBP2,NUP43,NUP35,VPS37B,NUP133 |
| Protein transport | GO:0015031 | 17 | 6.95x10^-7^ | | NUP160,COPE,UNC119,AP1M1,STX10,RANBP2,NUP43,MTX3,NUP35,VPS37B,COG6,NUP133,CCDC91,GPR89B,SLC15A3,CSE1L,SNX5 |
| Protein phosphorylation | GO:0006468 | 16 | 2.18x10^-6^ | | PRKD2,STK38L,MAP4K5,BCKDK,PRPF4B,CAMK2G,CAMK2D,PRKAB2,PAK2,MAP3K3,GSK3B,PDIK1L,FES,GRK2,RIOK2,TRIB2 |
| mRNA transport | GO:0051028 | 8 | 3.20x10^-6^ | | NUP160,NCBP2,QKI,SRSF7,RANBP2,NUP43,NUP35,NUP133 |
| **T2DM *vs* BC** | | | | | |
| Hemoglobin metabolic process | GO:0020027 | 2 | 3.38x10^-4^ | | EPB42,AHSP |
| Cell cycle arrest | GO:0007050 | 3 | 2.38x10^-3^ | | PPP1R15A,CDKN2D,STRADB |
| Peptide cross-linking | GO:0018149 | 2 | 4.23x10^-3^ | | CD3G,EPB42 |
| Protein transport | GO:0015031 | 4 | 6.12x10^-3^ | | CETN3,CD3G,SELENBP1,VPS37B |
| Positive regulation of endoplasmic reticulum stress-induced eif2 alpha dephosphorylation | GO:1903917 | 1 | 9.27x10^-3^ | | PPP1R15A |
| Transcription elongation from mitochondrial promoter | GO:0006392 | 1 | 9.27x10^-3^ | | TEFM |
| Regulation of lymphocyte apoptotic process | GO:0070228 | 1 | 9.27x10^-8^ | | CD3G |
| Positive regulation of translational initiation in response to stress | GO:0032058 | 1 | 9.27x10^-3^ | | PPP1R15A |
| Cell morphogenesis | GO:0000902 | 2 | 9.71x10^-3^ | | EPB42,STRADB |
| Protein adenylylation | GO:0018117 | 1 | 1.04x10^-2^ | | SELENOO |

**Table S2.** Differentially expressed genes (DEGs) annotated by the Cytoscape tool.

| **Comparison** | **Total DEGs** | **Annotated DEGs in Cytoscape** | **Included in Main PPI Network** | **Excluded from the PPI network** | **Additional Interactors** | **Hub** **Genes** |
| --- | --- | --- | --- | --- | --- | --- |
| T2DM vs PC | 113 | 97 | 35 | 62 | 30 | 8 |
| T2DM vs LC | 275 | 267 | 112 | 155 | 30 | 5 |
| T2DM vs BC | 26 | 25 | 9 | 16 | 30 | 9 |

**Table S3.** Top 20 ranked genes for T2DM *vs* PC and T2DM *vs* LC, shorlisted on the basis of 11 different topological features from PPI network analysis. The last row in each case indicate common hub genes which were found to be top ranked as well as common for all listed topological features.

| **Topological Feature** | **Top 20 genes ranked by score (obtained from cytoHubba)** |
| --- | --- |
| **T2DM *vs* PC** |  |
| Degree | CST3, TIMP1, P4HB, IGFBP3, CTSA, TRAPPC1, GRN, PYCARD, VAMP2, ITGB2, AP1M1, RAC1, PAK2, ATP11A, CD14, HERC4, RNF126, CFP, MGST1, REL |
| Maximal Clique Centrality | IGFBP3, P4HB, TIMP1, CST3, CTSA, TRAPPC1, GRN, PYCARD, VAMP2, ITGB2, AP1M1, ATP11A, RAC1, PAK2, HERC4, RNF126, CFP, CD14, MGST1, REL |
| Density of Maximum Neighborhood Component | IGFBP3, CTSA, TRAPPC1, GRN, PYCARD, P4HB, TIMP1, CST3, ATP11A, PAK2, HERC4, RNF126, VAMP2, CFP, CD14, MGST1, REL, RAB3D, EME1, AGTRAP |
| Maximum Neighborhood Component | CST3, TIMP1, P4HB, IGFBP3, CTSA, TRAPPC1, GRN, PYCARD, VAMP2, ITGB2, AP1M1, RAC1, ATP11A, PAK2, HERC4, RNF126, CFP, CD14, MGST1, REL |
| Edge Percolated Component | CST3, TIMP1, P4HB, IGFBP3, TRAPPC1, GRN, PYCARD, CTSA, VAMP2, ITGB2, AP1M1, RAC1, PAK2, ATP11A, CFP, AGTRAP, REL, CD14, COL18A1, RNF126 |
| Bottleneck | TIMP1, RAC1, CST3, VAMP2, BSG, CEBPD, P4HB, IGFBP3, TRAPPC1, GRN, PYCARD, CTSA, ITGB2, AP1M1, PAK2, ATP11A, CFP, AGTRAP, REL, CD14 |
| EcCentricity | RAC1, ITGB2, TIMP1, CST3, BSG, P4HB, IGFBP3, TRAPPC1, GRN, PYCARD, CTSA, AP1M1, PAK2, ATP11A, AGTRAP, REL, CD14, COL18A1, BRK1, MCL1 |
| Closeness | CST3, TIMP1, P4HB, IGFBP3, ITGB2, VAMP2, TRAPPC1, GRN, PYCARD, CTSA, RAC1, AP1M1, PAK2, MGST1, ATP11A, CD14, REL, MCL1, BRK1, AGTRAP |
| Radiality | TIMP1, CST3, P4HB, ITGB2, IGFBP3, RAC1, VAMP2, TRAPPC1, GRN, PYCARD, CTSA, AP1M1, PAK2, MGST1, BRK1, ATP11A, CD14, MCL1, REL, AGTRAP |
| Betweenness | RAC1, ITGB2, BSG, CEBPD, TIMP1, PAK2, CST3, VAMP2, CD14, P4HB, AP1M1, REL, MGST1, COL18A1, CFP, IGFBP3, TRAPPC1, GRN, PYCARD, CTSA |
| Stress | ITGB2, RAC1, CEBPD, PAK2, TIMP1, BSG, CD14, VAMP2, CST3, P4HB, REL, AP1M1, MGST1, COL18A1, CFP, IGFBP3, TRAPPC1, GRN, PYCARD, CTSA |
| *Common hub genes* | *CST3, CTSA, GRN, IGFBP3, P4HB, PAK2, TIMP1, TRAPPC1* |
| **T2DM *vs* LC** | |
| Degree | NCBP2, RANBP2, NUP133, NUP160, NUP43, SRSF7, CRNKL1, PRPF31, NUP35, ZWILCH, MIS12, GSK3B, CDC23, UBE2Q2, HERC4, ZBTB16, RNF126, HACE1, POLK, ATR |
| Maximal Clique Centrality | PRPF31, CRNKL1, SRSF7, NCBP2, RANBP2, NUP133, NUP160, NUP43, NUP35, ZWILCH, MIS12, CDC23, UBE2Q2, HERC4, ZBTB16, RNF126, HACE1, GSK3B, POLK, ATR |
| Density of Maximum Neighborhood Component | PRPF31, CRNKL1, ZWILCH, MIS12, SRSF7, NUP35, UBE2Q2, HERC4, ZBTB16, RNF126, HACE1, INTS13, ICE2, CDC23, RANBP2, NUP133, NUP160, NUP43, NCBP2, MLLT3 |
| Maximum Neighborhood Component | NCBP2, RANBP2, NUP133, NUP160, NUP43, SRSF7, CRNKL1, PRPF31, NUP35, ZWILCH, MIS12, CDC23, GSK3B, UBE2Q2, HERC4, ZBTB16, RNF126, HACE1, POLK, ATR |
| Edge Percolated Component | NCBP2, NUP133, RANBP2, NUP160, NUP43, SRSF7, CRNKL1, PRPF31, NUP35, ZWILCH, MIS12, GSK3B, ATR, POLK, CDC23, UBE2Q2, INTS13, ICE2, HERC4, ZBTB16 |
| Bottleneck | RANBP2, QKI, TOLLIP, CFP, NCBP2, HDAC5, MED23, ALDOA, NUP133, PAK2, GSK3B, MTERF3, ABCE1, AP1M1, MSH2, WDR36, IFT80, CEBPA, NUP160, NUP43 |
| EcCentricity | RANBP2, NUP133, NUP160, NUP43, NCBP2, PAK2, GSK3B, ABCE1, AP1M1, MSH2, SRSF7, CRNKL1, PRPF31, NUP35, ZWILCH, MIS12, CDC23, GSPT2, PRPF39, TUBA4A |
| Closeness | NCBP2, RANBP2, NUP133, NUP160, NUP43, GSK3B, SRSF7, CDC23, PAK2, ZWILCH, MIS12, CRNKL1, PRPF31, NUP35, ATR, POLK, UBE2Q2, HERC4, ZBTB16, RNF126 |
| Radiality | RANBP2, NUP133, NUP160, NUP43, NCBP2, GSK3B, PAK2, CDC23, ZWILCH, MIS12, SRSF7, GSPT2, TUBA4A, ABCE1, NUP35, MSH2, ATR, POLK, UBE2Q2, HERC4 |
| Betweenness | QKI, TOLLIP, CFP, ALDOA, MED23, GSK3B, RANBP2, HDAC5, PAK2, NUP133, NUP160, NUP43, NCBP2, ABCE1, MTERF3, IFT80, CEBPA, WDR36, NOC2L, ITGAV |
| Stress | QKI, MED23, RANBP2, NCBP2, GSK3B, TOLLIP, HDAC5, CFP, NUP133, NUP160, NUP43, IFT80, MTERF3, ALDOA, PAK2, CEBPA, ATR, POLK, SRSF7, CDC23 |
| *Common hub genes* | *NCBP2, NUP133, NUP160, NUP43, RANBP2* |

**Table S4.** Description and function of identified common hub gene for T2DM *vs* PC, T2DM *vs* LC and T2DM *vs* BC.

| **Gene** | **Description** | **Function** |
| --- | --- | --- |
| **T2DM *vs* PC** | | |
| CST3 | Neuroendocrine basic polypeptide | Belongs to the cystatin family, which act as an inhibitor of cysteine proteinases. |
| CTSA | Protective protein for beta-galactosidase | These are essential for both the stability and activity of beta-galactosidase and neuraminidase. This protein is also a carboxypeptidase and can deamidate tachykinins. |
| GRN | Proepithelin | Granulins have possible cytokine-like activity. They may play a role in inflammation, wound repair, and tissue remodeling. |
| IGFBP3 | Insulin-like growth factor binding protein 3 | They alter the interaction of IGFs with their cell surface receptors. Also exhibits IGF-independent antiproliferative and apoptotic effects mediated by its receptor TMEM219/IGFBP-3R. |
| P4HB | Cellular thyroid hormone-binding protein | It catalyzes the formation, breakage and rearrangement of disulfide bonds, thus invloved in protein folding.. |
| PAK2 | P21 protein (Cdc42/Rac)-activated kinase 2 | It plays a role in a variety of different signaling pathways including cytoskeleton regulation, cell motility, cell cycle progression, apoptosis or proliferation. It also phosphorylates JUN and plays an important role in EGF-induced cell proliferation. |
| TIMP1 | Tissue inhibitor of metalloproteinases 1 | It irreversibly inactivates MMPs such as collgenases, by binding to their catalytic zinc cofactor. Also functions as a growth factor that regulates cell differentiation, migration and cell death and activates cellular signaling cascades via CD63 and ITGB1. Plays a role in integrin signaling. |
| TRAPPC1 | Trafficking protein particle complex subunit 1 | Belongs to the TRAPP small subunits family, BET5 subfamily. It may play a role in vesicular transport from endoplasmic reticulum to golgi.. |
| **T2DM *vs* LC** | | |
| NCBP2 | Nuclear cap binding protein subunit 2 | It is component of the cap-binding complex, which binds co-transcriptionally to the 5' cap of pre-mRNAs and is involved in various processes such as pre-mRNA splicing, translation regulation, nonsense-mediated mRNA decay, RNA-mediated gene silencing (RNAi) by miRNAs and mRNA export. |
| NUP133 | Nuclear pore complex protein 133 | It is a nucleoporin, involved in poly(A)+ RNA transport. |
| NUP160 | Nuclear pore complex protein 160 | It is involved in poly(A)+ RNA transport, mediates nucleoplasmic transport. |
| NUP43 | Nuclear pore complex protein 43 | It is a component of the Nup107-160 subcomplex of the nuclear pore complex (NPC). The Nup107-160 subcomplex is also required for kinetochore-microtubule attachment, mitotic progression and chromosome segregation. |
| RANBP2 | Nuclear pore complex protein 358 | It is a component of the nuclear export pathway. |
| **T2DM *vs* BC** | | |
| ALYREF | Transcriptional coactivator Aly/REF | It is an export adapter involved in nuclear export of spliced and unspliced mRNA. It is also involved in transcription elongation and genome stability. |
| CD3G | T-cell surface glycoprotein CD3 gamma chain | It is part of the TCR-CD3 complex present on T-lymphocyte cell surface that plays an essential role in adaptive immune response. |
| CENPK | Interphase centromere complex protein 37 | It is component of the CENPA-CAD (nucleosome distal) complex, a complex recruited to centromeres which is involved in assembly of kinetochore proteins, mitotic progression and chromosome segregation. |
| EFHD2 | EF-hand domain-containing protein D2 | It may regulate B-cell receptor (BCR)-induced immature and primary B-cell apoptosis. It plays a role as negative regulator of the canonical NF-kappa-B-activating branch. It also controls spontaneous apoptosis through the regulation of BCL2L1 abundance. |
| JUNB | Transcription factor jun-B | It is a transcription factor involved in regulating gene activity following the primary growth factor response. |
| PPP1R15A | Myeloid differentiation primary response protein MyD116 homolog | It recruits the serine/threonine-protein phosphatase PP1 to dephosphorylate the translation initiation factor eIF-2A/EIF2S1, thereby reversing the shut-off of protein synthesis initiated by stress-inducible kinases and facilitating recovery of cells from stress. It down-regulates the TGF-beta signaling pathway by promoting dephosphorylation of TGFB1 by PP1. It may promote apoptosis by inducing TP53 phosphorylation on 'Ser-15'. |
| UBE2M | Ubiquitin-conjugating enzyme E2 M | It accepts the ubiquitin-like protein NEDD8 from the UBA3- NAE1 E1 complex and catalyzes its covalent attachment to other proteins. |
| VPS37B | Vacuolar protein sorting 37 homolog B | It is a component of the ESCRT-I complex, a regulator of vesicular trafficking process. It is required for the sorting of endocytic ubiquitinated cargos into multivesicular bodies. It may be involved in cell growth and differentiation. |
| WWP2 | WW domain containing E3 ubiquitin protein ligase 2 | It is an E3 ubiquitin-protein ligase which accepts ubiquitin from an E2 ubiquitin-conjugating enzyme in the form of a thioester and then directly transfers the ubiquitin to targeted substrates. |
